# Visualizing the organization and differentiation of the male-specific nervous system of *C. elegans*

**DOI:** 10.1101/2021.04.06.438718

**Authors:** Tessa Tekieli, Eviatar Yemini, Amin Nejatbakhsh, Erdem Varol, Robert W. Fernandez, Neda Masoudi, Liam Paninski, Oliver Hobert

## Abstract

Sex differences in the brain are prevalent throughout the animal kingdom and particularly well appreciated in the nematode *C. elegans*. While 294 neurons are shared between the two sexes, the nervous system of the male contains an additional 93 male-specific neurons, most of which have received very little attention so far. To make these neurons amenable for future study, we describe here how a multicolor, multipromoter reporter transgene, NeuroPAL, is capable of visualizing the distinct identities of all male specific neurons. We used this tool to visualize and characterize a number of features of the male-specific nervous system. We provide several proofs of concept for using NeuroPAL to identify the sites of expression of *gfp-tagged* reporter genes. We demonstrate the usage of NeuroPAL for cellular fate analysis by analyzing the effect of removal of developmental patterning genes, including a HOX cluster gene (*egl-5*), a miRNA (*lin-4*) and a proneural gene (*lin-32/Ato*), on neuronal identity acquisition within the male-specific nervous system. We use NeuroPAL and its intrinsic cohort of more than 40 distinct differentiation markers to show that, even though male-specific neurons are generated throughout all four larval stages, they execute their terminal differentiation program in a coordinated manner in the fourth larval stage that is concomitant with male tale retraction. This wave of differentiation couples neuronal maturation programs with the appearance of sexual organs. We call this wave “just-in-time” differentiation by its analogy to the mechanism of “just-in-time” transcription of metabolic pathway genes.

## INTRODUCTION

It is generally appreciated that nervous systems are sexually dimorphic on a gross anatomical level. However, sex differences in nervous systems have been carefully mapped out, with single-cell resolution, in only very few animals. The nematode *C. elegans* is the only organism for which a complete cellular, lineage, and anatomical map of the entire nervous system has been described for both sexes (**Fig.1A,B, Fig.S1**)(Cook et al., 2019; Jarrell et al., 2012; Sulston et al., 1980; Sulston and Horvitz, 1977). With 383 neurons total, the nervous system of the male is almost 30% larger than that of the hermaphrodite (302 neurons). Based on lineage and anatomy and molecular profiles, 294 neurons are shared between both sexes. Hermaphrodites, which are somatic females, contain an additional 8 hermaphrodite-specific neurons that fall into two classes: the well characterized HSN and VC motor neuron classes, both of which control egg laying behavior (Schafer, 2005). The male contains an additional 93 neurons that fall into 27 anatomically distinct classes (**Fig.1A,B, Table S1**)(Cook et al., 2019; Molina-Garcia et al., 2020; Sammut et al., 2015; Sulston et al., 1980). These 27 neuron classes are extensively interconnected and the structure of their interconnectivity displays a number of notable features, including modular substructures regulating subsequences of male mating behavior; multiple, parallel and short synaptic pathways directly connecting sensory neurons to end organs and recurrent, reciprocal connectivity among the male’s many sensory neurons (Cook et al., 2019; Jarrell et al., 2012)

**Fig.1:**
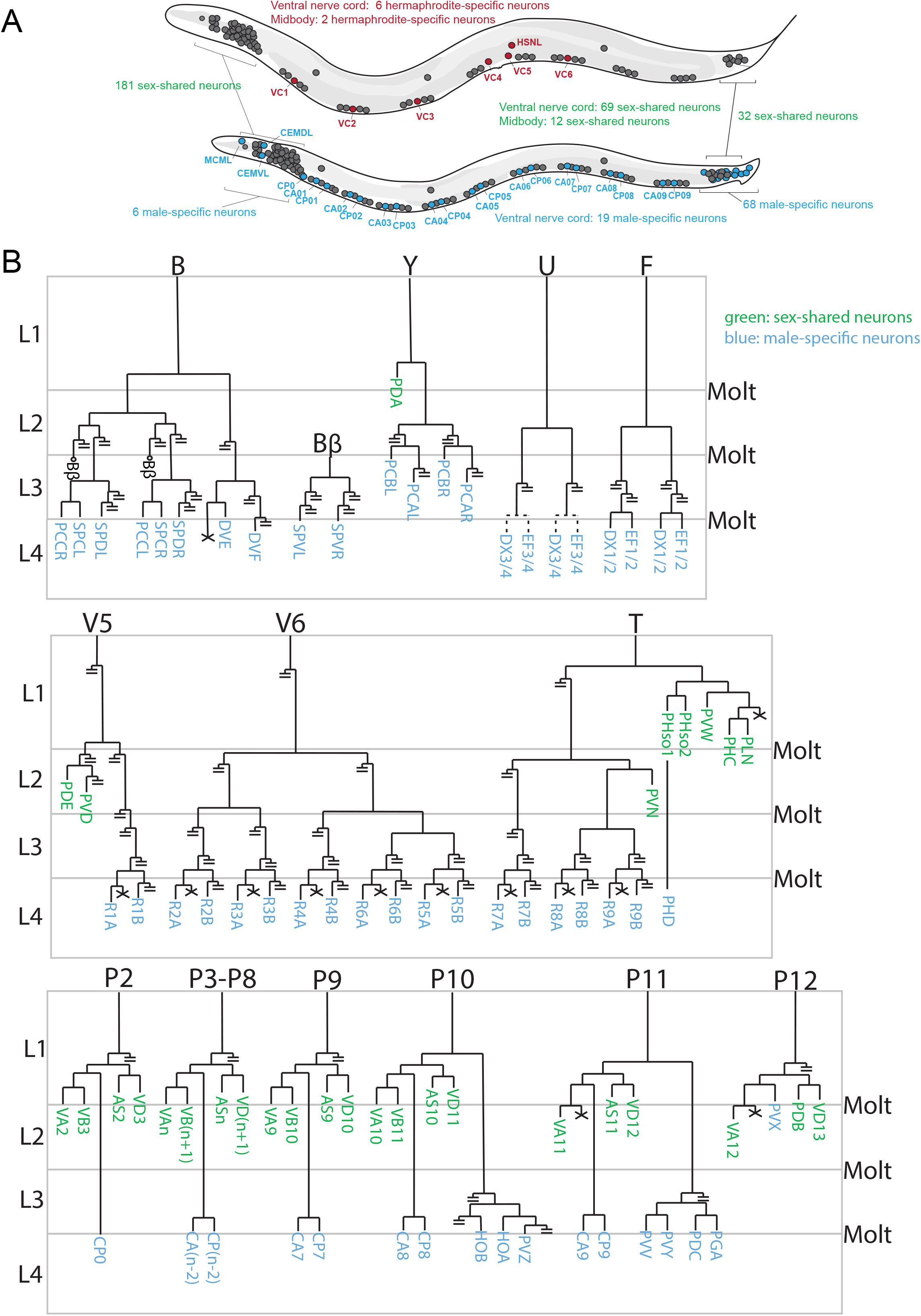
Lineage patterns to illustrate temporal generation. **A:** Schematic overview of sex-specific nervous systems. **B:** Abridged version of the Sulston (1980) lineage diagram of male-specific neurons in the tail. Only those branches that generate neurons are shown till the end, cells that die are denoted with an X, all other branches are cut off (with double strike through). Note that male-specific neurons are generated at different times of larval development, with the earliest being generated in the L1 stage (PVX). The lineage of the male-specific CEM neurons is not shown since it is generated within the embryo and dies in hermaphrodites.

Of the 27 male-specific neuron classes, two are the head sensory neuron classes CEM and MCM, two are the ventral nerve cord motor neuron classes CA and CP, and the remaining 23 classes are located in the tail of the animal. Some of the 27 male-specific neuron classes are composed of only a single neuron or two bilaterally symmetric neurons. Other neuron classes are composed of multiple class members: the A- and B-type ray sensory neurons are each composed of nine distinct bilateral pairs. With the exception of the four CEM sensory neurons in the head, which are born in the embryo and induced to die in hermaphrodites, all male-specific neurons are generated during postembryonic development from blast cells that proliferate and differentiate in a male-specific manner (**Fig.1B**)(Sulston et al., 1980). Based on cell divisions patterns, the 87 postembryonically generated male-specific neurons are generated at different larval stages. Each individual larval stage contributes to the generation of some of these postembryonic neurons (Sulston et al., 1980). However, when exactly these neurons terminally differentiate is poorly understood. Moreover, in his classic lineage studies Sulston also noted that the number of two male-specific neuron classes, DX and EF, display a variable number of class members (Sulston et al., 1980). Since this observation was originally based on Nomarski optics and limited sample size, this variability has not been well characterized and has not been observed elsewhere within or outside the nervous system of *C. elegans*.

The vast majority of the sex-shared nervous system is generated in the embryo and synaptically connected by the first larval stage. Thus, one fascinating problem presented by the male-specific nervous system is how the many postembryonically generated, male-specific neurons become integrated into already existing circuitry. Of the 27 male-specific neuron classes, all but one (PCC) make synaptic contacts to sex-shared neurons. Understanding how such integration occurs may provide interesting insights for more complex vertebrate nervous systems, which are similarly characterized by the addition of new neurons throughout many stages of juvenile and even adult stages.

Despite many interesting aspects of the male nervous system, it has received little attention over the years when compared to the nervous system of the hermaphrodite. A number of studies have illuminated aspects of the development and function of male-specific neurons, but those studies only dealt with a limited set of neurons (Barr et al., 2018; Emmons, 2014, 2018; Garcia et al., 2001; Garcia and Portman, 2016; Liu and Sternberg, 1995; Portman, 2017). Hence, many aspects of the development and function of the 93 male-specific neurons remain uncharted territory. With some notable exceptions, including the systematic mapping of neurotransmitter identities (Gendrel et al., 2016; Pereira et al., 2015; Serrano-Saiz et al., 2017), marker analysis in the ray sensory neurons (Lints et al., 2004) and ventral nerve cord (Kalis et al., 2014), few molecular markers have been developed that label male-specific neurons. Single-cell transcriptome approaches have so far exclusively focused on the hermaphrodite (Cao et al., 2017; Packer et al., 2019; Taylor et al., 2020). This dearth of molecular markers not only limits the ability to assess, for example, cell fate in specific mutant backgrounds, but also complicates the means by which cellular expression patterns in the male tail can be unambiguously identified.

Here, we address these shortcomings by showing that NeuroPAL, a previously described multicolor transgene that distinguishes all neuron classes in the hermaphrodites (Yemini et al., 2021), can also be used to disambiguate the 93 neurons of the male nervous system. We find that the NeuroPAL transgene, which harbors more than 40 promoters that drive the expression of four distinct fluorophores, generates a color map that provides sufficient discriminatory power to reliably identify all male-specific neurons. We provide proof-of-principle examples that show how to use NeuroPAL to identify gene expression patterns in the nervous system, and use the NeuroPAL color map to provide a number of insights into the development of the male-specific nervous system.

## RESULTS

### NeuroPAL provides discriminatory color barcodes for all male-specific neurons

With the exception of neurotransmitter pathway genes (Gendrel et al., 2016; Lints and Emmons, 1999; Pereira et al., 2015; Serrano-Saiz et al., 2017), few molecular markers have been comprehensively described for male-specific neurons (www.wormbase.org). For several related neuron classes, for example the ray neurons, molecular markers are available, but they do not provide sufficient resolution to distinguish between all individual class members (Lints and Emmons, 1999; Lints et al., 2004). We set out to test whether the NeuroPAL transgene that we previously described for the *C. elegans* hermaphrodite (Yemini et al., 2021) would provide a similarly information rich molecular map of the male-specific nervous system.

The NeuroPAL transgene was designed to provide color codes to all neurons of the *C. elegans* hermaphrodite (Yemini et al., 2021). This was achieved through the judicious use of four fluorophores with separable emission spectra (mTagBFP2, CyOFP1, Tag-RFP-T, mNeptune2.5), expressed under the control of a set of 43 different promoters with overlapping expression profiles (39 neuron-type specific promoters + 4 distinct, but fused panneuronal promoters) (Yemini et al., 2021). Promoter choices were dictated by the goal of having neighboring neurons display distinct color codes, thereby unambiguously discriminating neighboring neuron identities from one another.

We found that three NeuroPAL transgenes (independently integrated transgenes *otIs696, otIs669* or *otIs670*) distinguish all neighboring male-specific neurons from one another. This is illustrated in the whole-animal overview panel in **Fig.2A** along with large-scale images of all regions of the *C. elegans* nervous system that contain male-specific neurons (**Fig.2B-E**). The origin of the color code for each neuron is listed in **Table S1**. NeuroPAL not only provides color codes for neuron classes for which few or no molecular markers were previously available, but it also distinguishes neuronal subclasses that could previously not be discriminated. For example, each individual subclass of A- and B-type motor neurons has its own unique color code (**Fig.2, Table S1**).

**Fig.2:**
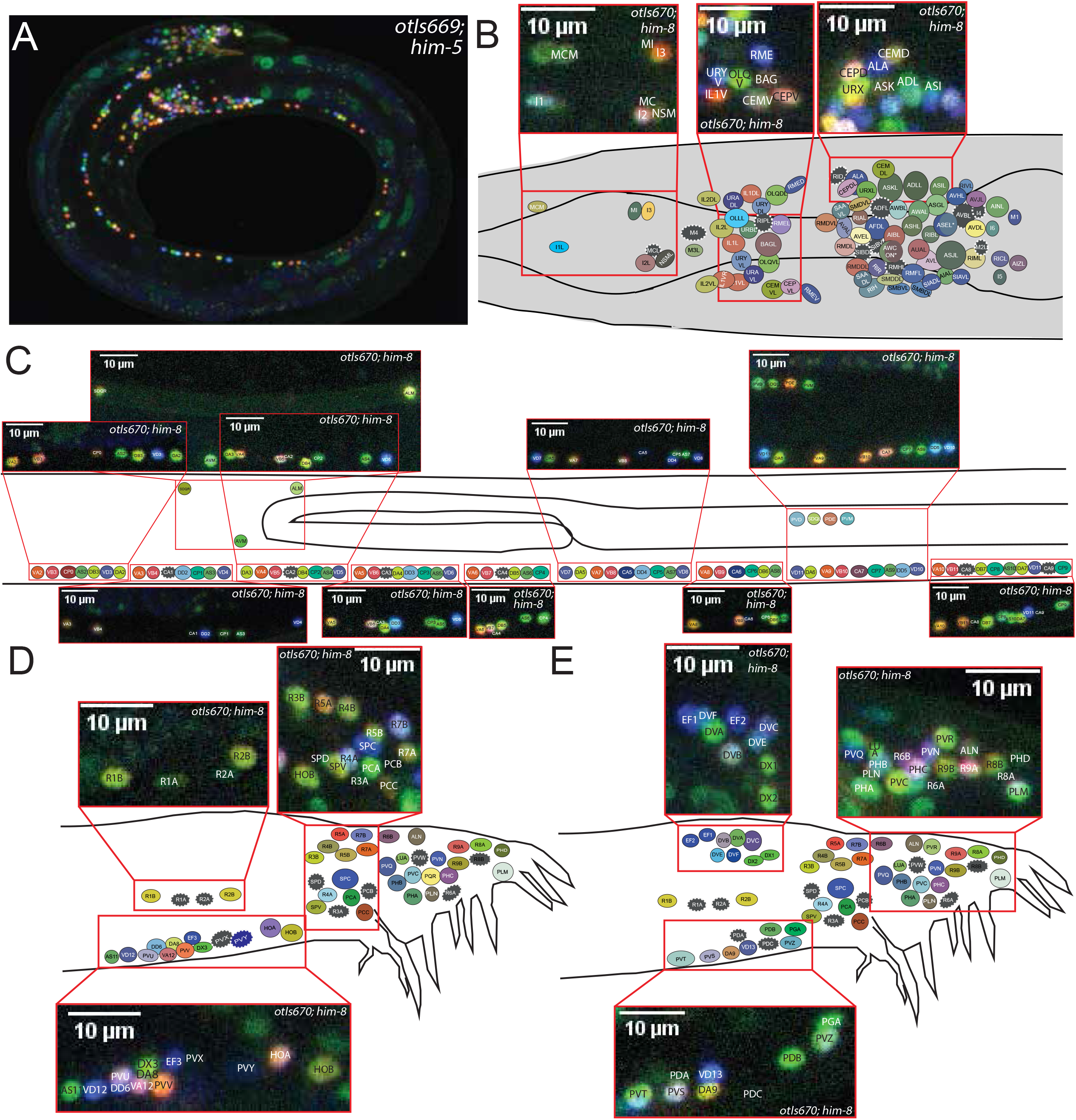
Multicolor map of the adult, male-specific nervous system. **A:** Overview of entire male NeuroPAL worm (*otIs669;him-5*). **B:** Schematic NeuroPAL map of the male head with images containing male-specific neurons shown above the schematic. The leftmost image depicts the anterior pharynx containing the male-specific MCM neurons. The middle image depicts the anterior ganglion containing the-male specific CEMV neurons. The rightmost image depicts the dorsal ganglion containing the male-specific CEMD neurons. **C:** Schematic NeuroPAL map of the male ventral nerve cord (VNC). Real images of the VNC from adult male worms are depicted above. **D:** Schematic NeuroPAL map of the left side of the male tail. Real images, from the indicated tail regions, depict the neurons that are contained within those regions. **E:** Schematic NeuroPAL map of the right side of the male tail. Real images, from the indicated tail regions, depict the neurons that are contained within those regions.

### Using NeuroPAL to address stereotypy in the male-specific nervous system

We first used NeuroPAL to address questions that relate to stereotypy of the male-specific nervous system. In his original lineage analysis of the male tail, John Sulston reported on an unusual phenomenon, not observed anywhere else in the entire organism: descendants of the U ectoblast produce variable numbers of DX and EF neurons, a notion indicated by stippled lines in Sulston’s original lineage diagram (redrawn here in **Fig.1B**). This violates the complete stereotypy and deterministic nature of all cell lineages, both neuronal and non-neuronal. Moreover, according to the Sulston lineage diagram, this variability is restricted to the EF and DX neurons that descend from the U neuroblast and that are located in the preanal ganglion (the EF3 & 4 and DX3 & 4 neurons). In contrast, the DX and EF neurons that are produced from the F neuroblast (EF1 & 2, DX1 & 2), located in the dorsal rectal ganglion, were generated in an apparently invariant manner (as per the Sulston lineage diagram) (**Fig.1B**). However, no quantification of this was provided. Because the lineage analysis entirely relied on cleavage pattern alone, it was also not clear to what extent the variably produced DX and EF neurons acquire a differentiated state.

Using NeuroPAL, we examined 22 young adult males and found variability in the presence of fully differentiated EF and DX neurons in the preanal ganglion (**Fig.3A**) – assessed by wild-type expression of NeuroPAL colors in these neurons. Within the F-derived dorsorectal ganglion, 22/22 animals invariably showed two fully differentiated DX neurons (DX1 and DX2) and two EF neurons (EF1 and EF2), corroborating John Sulston’s observations. In the U-derived preanal ganglion, 19/22 animals show one DX and one EF neuron (= DX3 and EF3), 1/22 had one additional EF (= EF4), and 2/22 had one additional EF (= EF4) and one additional DX (= DX4).

**Fig.3:**
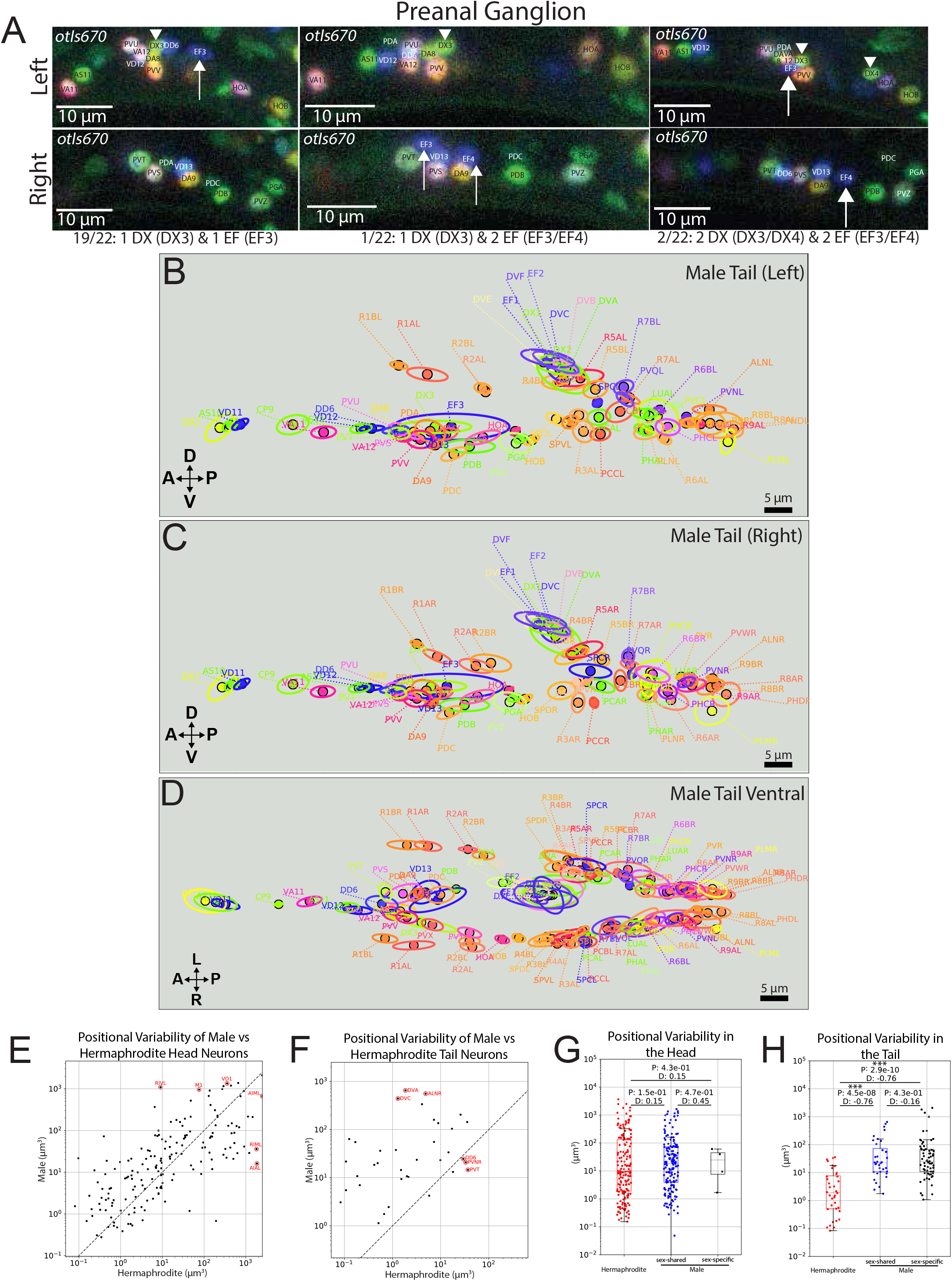
Variability of cell generation and position in the adult male tail. **A:** NeuroPAL (*otIs670; him-8*) is used to visualize the variability of DX and EF neuron (DX3/4 and EF3/4) generation in the preanal ganglion of the adult male tail. Example images of the male preanal ganglion are shown in NeuroPAL worms. These images depict real observations of the indicated number of DX and EF neurons. The left side of the preanal ganglion is shown in the top set of images and the right side of the preanal ganglion is shown in the bottom set of images. The fraction of NeuroPAL worms observed, with the indicated DX/EF neuron count, is listed below the set of images depicting the indicated numbers of DX and EF neurons. Arrows denote the EF neurons and arrowheads denote the DX neurons in the NeuroPAL images. **B-D:** The atlas of male tail neuron positional variability (based on 13 male tails) for the left (B), right (C), and ventral (D) sided views. Dots indicate the mean position of each neuron. Ellipses indicate the positional variability of each neuron in the given axis. Neurons colors approximate those in NeuroPAL but have been brightened for visibility. **E-F:** Positional variability of the individual hermaphrodite versus male neurons in the head (E) and tail (F). Six neurons that show maximal differences between both sexes are circled and identified. **G-H:** Quantification of positional variability for the collection of all head (G) and tail (H) neurons of the hermaphrodite (which are all sex-shared) versus the male sex-shared and sex-specific neurons. In the head, the positional variability is nearly the same for these three neuron groups. In the tail, the positional variability for the group of hermaphrodite neurons is far less than that of the male sex-shared and sex-specific neurons. We report the P-value (Mann-Whitney U test) for differences between hermaphrodites and males and the effect size (Cohen’s D). For the head N = 10 hermaphrodites, 12 males, 182 sex-shared neurons, and 6 male-specific neurons, with a mean of 9.6 neurons/hermaphrodite and 9.8 neurons/male. For the tail N = 10 hermaphrodites, 13 males, 41 sex-shared neurons, and 69 male-specific neurons, with a mean of 9.6 neurons/hermaphrodite and 11.6 neurons/male. Further hermaphrodite and male atlases can be found in Suppl. Fig.S1. Data is available in Suppl. Table S2.

The EF and DX neurons are also the neurons with the greatest inter-animal variability in their relative positioning. We arrived at this concluision by closely considering the overall variability of positioning of both sex-shared, as well as sexspecific neurons in the tail of the animal. We had previously shown that in the hermaphrodite head, where the vast majority of neurons are generated embryonically, most cells are positioned within a small volume of variability (Yemini et al., 2021) and we observer a similar extent of variability in the male head (**Fig.3**; **Table S2**). However, in the tail, where the vast majority of the postembryonically added male-specific neurons are located, there is substantially more positional variability, both in the sex-shared neurons as well as in the sex-specific neurons (**Fig.3; Table S2**). The EF and DX neurons stand out in the extent of variability in their positioning. It will be interesting to investigate whether the inter-animal variability in neuronal soma position in the male tail also translates into variability in neuronal process adjacency, and hence connectivity, between individual animals.

### Using NeuroPAL to characterize reporter gene expression patterns in the male tail

Compared to the hermaphrodite nervous system, there has been a remarkable scarcity of molecular markers for neurons in the male tail. The vast majority of reporter transgenes that researchers generate to analyze the expression of their gene of interest are usually only examined in hermaphrodites. One reason for the reluctance of identifying sites of reporter gene expression in the male tail has been the absence of reliable landmark reporters for most male-specific neurons. The fluorescence emission properties of NeuroPAL are designed to be separable from those of GFP signals. This allows researchers to overlay a GFP signal from a reporter gene of interest onto the neuron-specific color barcodes of NeuroPAL, thus identifying the sites of expression of GFP (or CFP/YFP)-based reporter transgenes. As a proof of principle, we analyzed *gfp*-tagged alleles of two neuropeptide encoding genes that were previously uncharacterized in males (*flp-3, flp-27*). To do so, we crossed these *gfp*-tagged alleles into a NeuroPAL background. We found that *flp-3::T2A::3xNLS::gfp* was expressed in the male-specific CA1, CA2, CA3, and CA4 neurons, located in the ventral nerve cord, as well as the male-specific R4A and SPV neurons in the tail (**Fig.4A)**. Using NeuroPAL for cell identification, we found that *flp-27::T2A::3xNLS::gfp* was expressed in the male-specific neurons CEMV, CEMD, CA8, CA9, PGA, R7A, and was dimly and variably expressed in R6B as well as the sex-shared neuron ASG **(Fig.4B**). These expression patterns corroborate the molecular diversity of members of the CA-type ventral nerve cord motor neurons, and that of the ray sensory neurons, as noted previously with other markers (Kalis et al., 2014; Lints et al., 2004).

**Fig.4:**
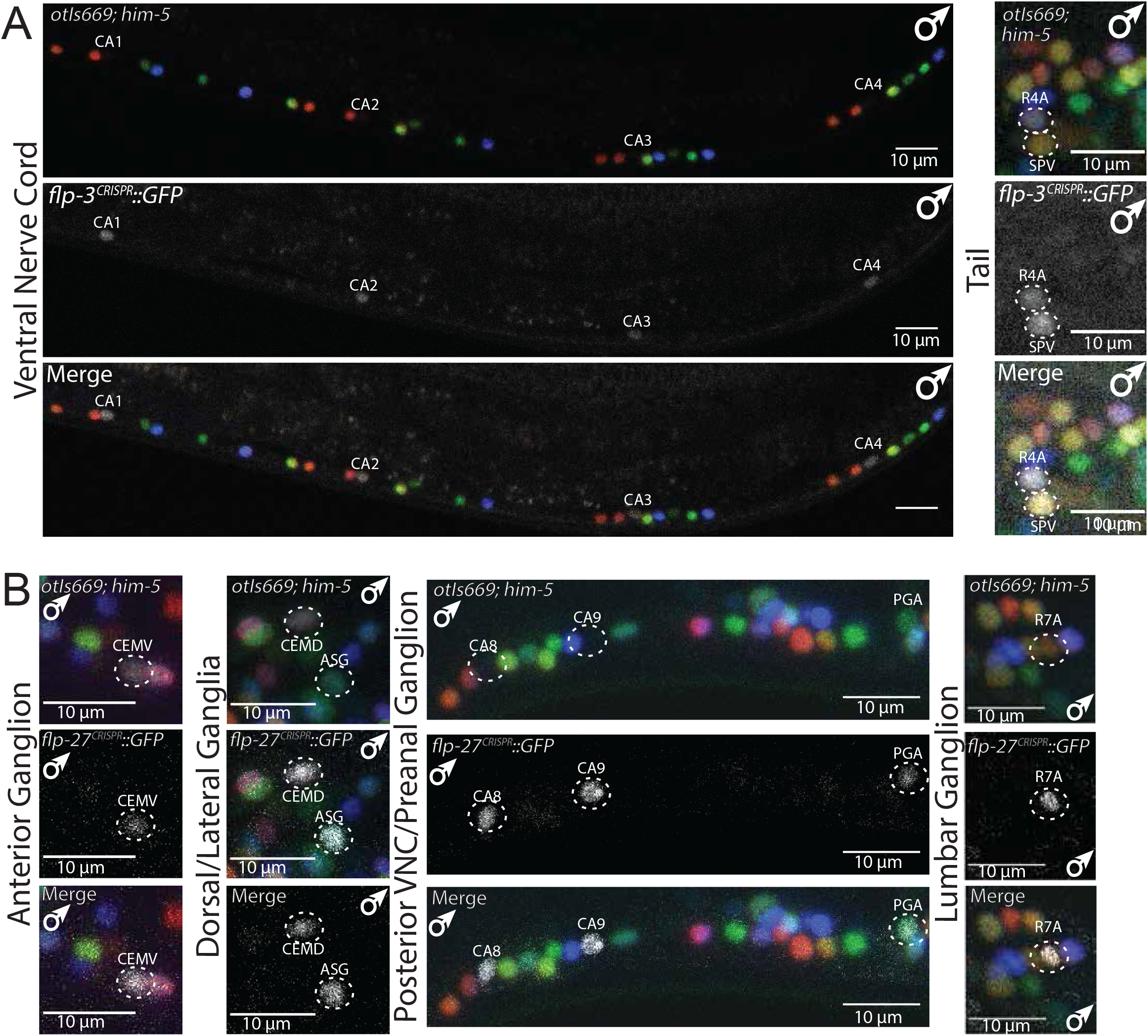
Using NeuroPAL for identification of *gfp-based* expression patterns. **A:** NeuroPAL (*otIs670; him-8*) is used to identify the GFP expression profile of a neuropeptide gene, *flp-3*, in the male ventral nerve cord (VNC) and the tail. In the VNC, *flp-3::GFP* is expressed in the male-specific neurons CA1-CA4, which are marked by labels on the image. In the tail, *flp-3::GFP* is expressed in the male-specific neurons R4A and SPV. **B:** NeuroPAL is used to identify the GFP expression profile of a neuropeptide gene, *flp-27*, in the male head, posterior VNC, preanal ganglion and lumbar ganglion. In the head, *flp-27* is expressed in the male-specific CEMV and CEMD neurons and in the sex-shared ASG neuron. In the posterior VNC, *flp-27* is expressed in the male-specific neurons CA8 and CA9, and in the male-specific neuron PGA in the preanal ganglion. In the lumbar ganglion, *flp-27* is expressed in the male-specific ray neuron R7A, and it is variably dimly expressed (not pictured) in R6B.

### Using NeuroPAL to measure neuronal cell-fate specification in the male tail

As described above, NeuroPAL is an indicator of expression for 39 neuron-type specific genes, as well as 4 panneuronal genes (that are fused together in the “UPN” construct), marking all male-specific neurons. These reporter genes measure a wide variety of phenotypic features of a neuron, including neurotransmitter synthesis and transport, neurotransmitter receptors, neuropeptides, sensory receptors from various families, and even panneuronal features (**Table S1**)(Yemini et al., 2021). The markers therefore provide a panoramic view of the differentiated state of all individual neurons, and this state can be probed for proper execution in mutant backgrounds.

We illustrate the NeuroPAL’s utility for such mutant analysis using three prominent patterning genes: a miRNA (*lin-4*)(Lee et al., 1993), a HOX cluster gene (*egl-5*)(Chow and Emmons, 1994), and a proneural bHLH gene (*lin-32/Atonal*)(Zhao and Emmons, 1995). The functions of these genes have been reported for only select parts of the male-specific nervous system and a comprehensive view of their function was therefore missing (Chalfie et al., 1981; Chow and Emmons, 1994; Zhao and Emmons, 1995). We sought to assess whether their reported defects can be recapitulated and better characterized with NeuroPAL. Furthermore, we anticipated identifying novel defects in these mutants in previously unexamined parts of the male-specific nervous system. Both of these expectations were fulfilled in all three cases examined, as described in the ensuing sections.

### NeuroPAL confirms predicted neuron losses and duplications in *lin-4/miRNA* mutants and identifies additional neuronal defects

Animals lacking the *lin-4* miRNA display an iteration of cellular fates normally executed in the first larval stage (Chalfie et al., 1981; Lee et al., 1993). This notion derives mainly from the analysis of the ectodermal V and T ectoblasts in the male. Specifically, V and T-derived ray neurons, normally generated in late larval stages, do not appear to be generated, based on neuronal nuclear morphology. On the other hand, L1-stage specific T-cell neurons appear to be duplicated, again based on nuclear morphologies (Chalfie et al., 1981)(redrawn in **Fig.5A**). We extended these previous findings through our ability to visualize neuronal differentiation programs with greater detail using NeuroPAL. We confirmed that ray neurons are not generated in *lin-4* mutant animals while neurons displaying the color code of the T-cell derived PHC, PHD, PLN, and PVW appear to be duplicated (**Fig.5B**).

**Fig.5:**
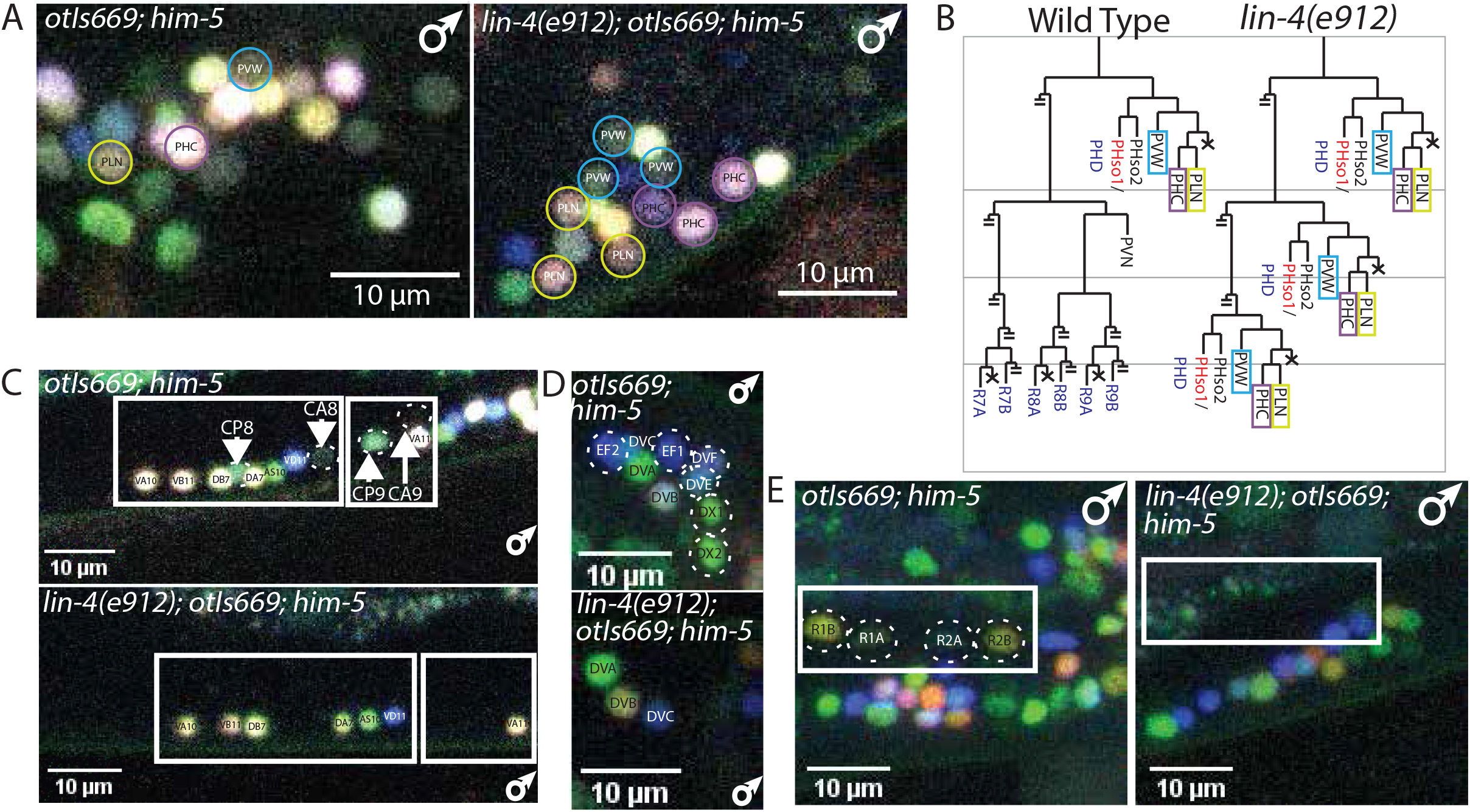
NeuroPAL visualizes neuronal-patterning defects in miRNA *lin-4* mutants. **A:** Lineage iterations in the L1-specific T lineage, first reported by Chalfie *et al*., 1981, are confirmed with NeuroPAL, as evidenced by the presence of additional neurons with the distinct NeuroPAL color barcodes of PLN, PHC, and PVW in the tail of *lin-4(e912)* males. The left image depicts a male NeuroPAL tail with a single PLN neuron (yellow circle), a single PHC neuron (purple circle), and a single PVW neuron (blue circle). The *lin-4(e912)* male tail contains three PLN neurons (yellow circles), three PHC neurons (purple circles), and three PVW neurons (blue circles), all distinguished by their NeuroPAL coloring. 20 animals were scored and all animals exhibited reiterations in the L1-specific T lineage. **B:** Lineage diagrams depicting the wild-type T lineage on the left and the *lin-4* mutant T lineage exhibiting lineage reiterations in the L1-specific T lineage, as reported by Chalfie *et al*., 1981. Only those lineage branches containing neurons are depicted to the end, cells that die are represented with an X, and all other branches are cut off (depicted by double strike through). Sex-shared neurons are depicted in black, hermaphroditespecific neurons are depicted in red, and male-specific neurons are depicted in blue. **C:** The color codes of the L3-specific CA8-9 and CP8-9 neurons in the male VNC are gone in *lin-4(e912)* mutants, while the L1-specific neurons VA, VB, VD are unaffected, as evidenced by the expression of their proper NeuroPAL colors. In the 20 animals that were scored, the color codes of the CA8 and CP8 neurons were never observed, while the color code of CA9 was observed in one animal, and the color code of CP9 was observed in five animals. **D:** Male-specific neurons generated from the B and F lineages in the dorsorectal ganglion (DRG), indicated by dashed circles in the image of NeuroPAL on its own, are gone in *lin-4(e912)* mutants. In contrast, sex-shared neurons (DVA, DVB, and DVC) remain in *lin-4(e912)* mutants. 20 animals were scored and neurons from the B and F lineages in the DRG were never observed. **E:** Male-specific ray neurons (R1A, R1B, R2A, R2B) generated from the V5 lineage, and depicted in dashed circles, are gone in *lin-4(e912)* mutants. Male-specific ray neurons were never observed in the 20 scored worms.

Cleavage defects in other lineages that produce male-specific neurons (B, Y, U, F) had been noted in *lin-4* mutant males (Chalfie et al., 1981), but whether and to what extent neuronal fate specification was disrupted in these lineages remained unclear. We observed no cells with color codes representative of the B, Y and F-derived neurons, suggesting that these neurons are not properly generated. This is consistent with a predicted “juvenizing” effect of *lin-4* mutants in the V and T-lineages, wherein earlier lineage patterns are re-iterated on the expense of neurons that are normally born at later larval stages. The color patterns in the preanal ganglion, where U cell descendants are normally located, was too complex to interpret, and therefore we cannot conclude whether there are defects in this lineage as well.

We also found that the fate of all male-specific neurons that are generated by the P neuroblast, are lost in *lin-4* mutant animals. In the tail, male-specific neuronal cell fates derived from the P10 and P11 lineages (HOA, HOB, etc.) appear to be lost. Lastly, in the ventral nerve cord, we observed a loss of color codes of the CA8-9 and CP8-9 neurons (**Fig.5C**). These neurons are normally generated by a cell division event in the L3 stage, and this division is possibly absent in *lin-4* mutants. In contrast, P-cell derived ventral nerve cord motor neurons, that are generated in both sexes in early larval stages, differentiate normally in *lin-4* null mutants (**Fig.5C**).

### NeuroPAL confirms the proneural function of *lin-32/Ato* in ray lineages and reveals additional proneural function in other lineages

The single ortholog of the proneural Atonal gene in *C. elegans, lin-32*, was initially identified and characterized based on its proneural role in the V5 and V6 ectoblast-derived ray neuron (Zhao and Emmons, 1995). Using NeuroPAL, we confirmed the loss of neuronal-fate specification in the ray lineage in *lin-32(tm1446)* mutants in the V5- and V6-derived ray neurons. This was evidenced by a lack of all color codes in these neurons, including the panneuronal color (**Fig.6**). The most posterior ray neurons, generated by the T-lineage (**Fig.1**), are unaffected in *lin-32* null mutants, as are all of the other T-derived neurons (**Fig.6**). Y, U and F-derived neurons, as well as P cell-derived, male-specific VNC motor neurons (see lineage diagram in **Fig.1**) are also unaffected in *lin-32* mutants (**Fig.6**). However, within the B lineage we discovered a novel proneural role of *lin-32* mutants. While the neurons generated by the posterior daughter of the B ectoblast cell (DVE and DVF, located in the dorsorectal ganglion) differentiate normally, the neuronal cell fates generated by the anterior daughter of B (mostly spicule neurons) cannot be detected (**Fig.6**).

**Fig.6:**
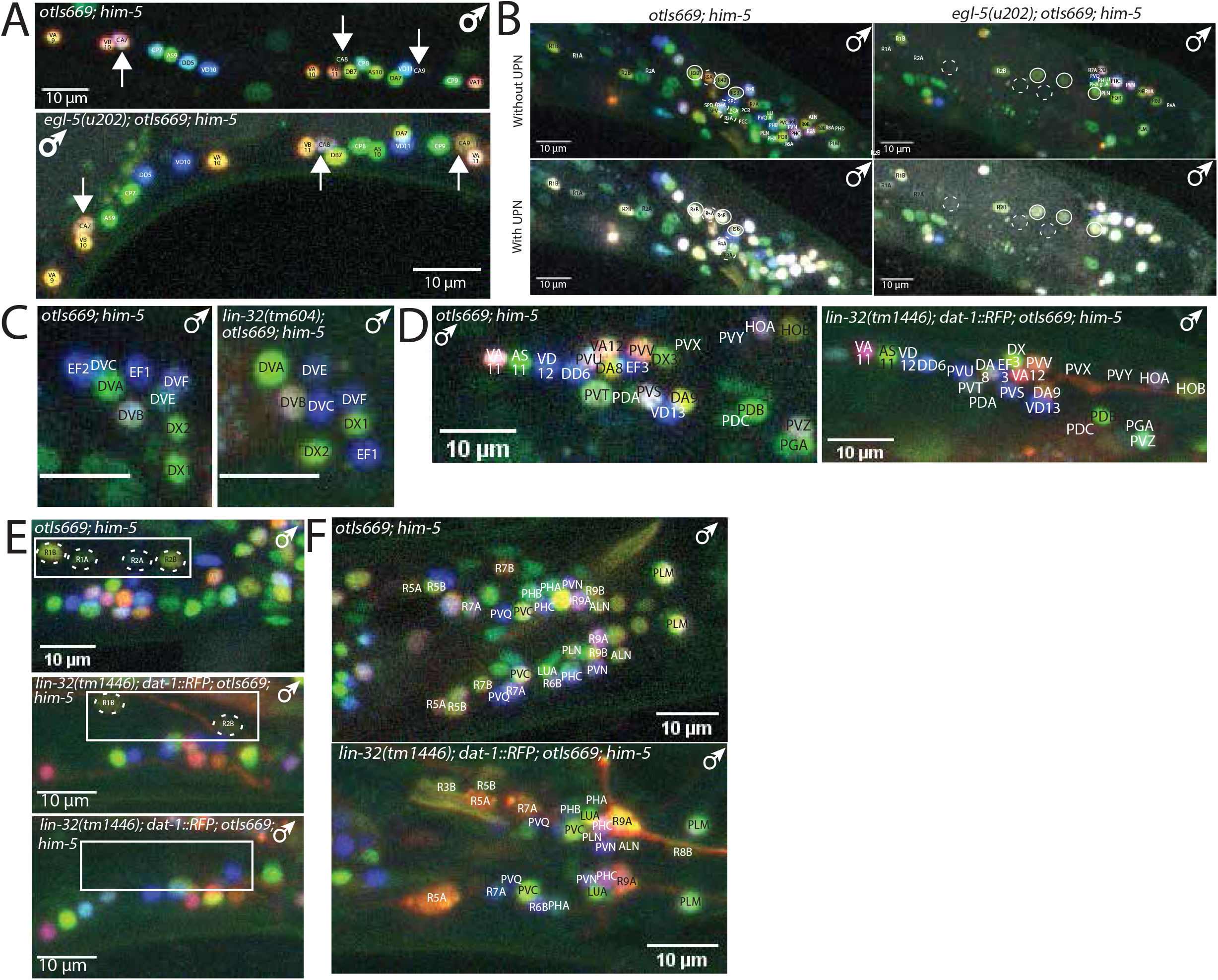
NeuroPAL visualizes patterning defects in transcription factor mutants. **A-B:** Patterning defects in the VNC of *egl-5* Hox mutant males are visualized using NeuroPAL (*otIs669; him-8*). **A:** The arrows denote CA7, CA8, and CA9 neurons in NeuroPAL and *egl-5(u202)*; NeuroPAL males. CA8 and CA9, normally marked only by the panneuronal marker in NeuroPAL, take on a distinct color that is similar to CA7 in *egl-5(u202)* males. The transformation of CA8 and CA9 neurons was found in all 14 animals that were scored. **B:** Transformations of ray neurons, R3, R4, and R5 to that of a R2 neuron fate was described by Lints *et al*., 2004. Here the A-type neurons (R3A, R4A, R5A) are denoted by dashed circles and the B-type neurons (R3B, R4B, R5B) are denoted by closed circles. In *egl-5*(*u202*) mutant males there is a loss of the NeuroPAL colors of the A- and B-type R3, R4, and R5 neurons and a gain of 3 R2A and 3 R2B neurons. The transformation of R3, R4, and R5 was observed in all 14 animals that were scored. **C-F:** Patterning defects in *lin-32*(*tm1446*) bHLH mutant animals, visualized with *otIs669*. **C:** All neurons in the dorsorectal ganglion (DRG) are unaffected in *lin-32(tm1446)* males as evidenced by the preservation of all NeuroPAL colors. Out of the 15 animals scored none of the *lin-32* mutant males showed defects in the DRG. **D:** All neurons in the preanal ganglion (PAG) are unaffected in *lin-32(tm1446)* males as evidenced by the preservation of all NeuroPAL colors. Out of the 15 animals scored none of the *lin-32* mutant males showed defects in the PAG. **E:** A variable loss of ray neurons R1B and R2B was observed in *lin-32(tm1446)* males. Two example images from *lin-32* mutant males are shown under a NeuroPAL image of the region of interest. In the middle *lin-32(tm1446)* image, R1B and R2B are present and properly colored, while in the bottom *lin-32(tm1446)* image both R1B and R2B are gone, thus demonstrating their variability in *lin-32* mutant males. Out of the 15 animals scored, nine had neither R1B nor R2B, three had one R2B, and three had both R1B and R2B. **F:** A representative image of a *lin-32(tm1446)* male depicts variable loss of male ray neurons in the tail. While R9A and R5A are observed in most *lin-32* mutant male tails, other ray neurons are variably identifiable by their NeuroPAL color. Out of the 15 *lin-32(tm1446)* males that were scored, all showed some defects in ray neurons. Within the same animal the ray neuron often differed between the left and right side as pictured.

### NeuroPAL identifies novel homeotic identity transformations in *egl-5* mutants

The B, Y, U and F ectoblasts, which divide exclusively in males, express the HOX cluster protein EGL-5 (Ferreira et al., 1999). In *egl-5* mutants, these ectoblasts fail to undergo proper divisions and generate no neurons (Chisholm, 1991). This conclusion was based on light-microscopy criteria: the absence of characteristic dense neuronal nuclei. Using the neuronal cell-fate markers present in the NeuroPAL transgene, we further corroborated this notion: none of the colored neurons that descend from B, Y, U and F (**Fig.1**) can be observed in *egl-5* mutants (**Fig.6**).

Previous work from the Emmons lab has revealed an anterior-posterior patterning role of the HOX genes *mab-5* and *egl-5* in the ray lineage (Emmons, 2005). Ray neuron 2 is *mab-5-*positive and *egl-5-*negative, while ray neurons 3 to 6 are *egl-5-* positive (Lints et al., 2004). It was suggested that in *egl-5* mutants, rays 3, 4, and 5 homeotically transform to the fate of ray 2 neurons. With the limited markers available at the time, this suggestion remained tentative (Lints et al., 2004). We verified this suggestion by confirming that color code changes in NeuroPAL are consistent with a ray 2 neuron identity transformation (**Fig.6**)

Using NeuroPAL, we discovered an additional homeotic identity transformation in the posterior-most, male-specific CA motor neurons (**Fig.6**). We found that the CA8 and CA9 neurons adopt the color code of the more anterior CA7 neuron. These transformations are conceptually similar to the homeotic transformations observed in the sex-shared, posterior-most DA and AS neurons in *egl-5* mutants (Kratsios et al., 2017). In those cases, the most posterior class member also transforms its identity to that of the more anteriorly located neuron.

### NeuroPAL reveals a coordinated differentiation wave that is concomitant with male tail retraction

We next used NeuroPAL as a tool to provide a panoramic view of timing of neuronal differentiation in the male tail. As illustrated in **Fig.1B**, male-specific neurons are generated at multiple distinct developmental stages. Some male-specific neurons are generated in the embryo (CEM neurons), more are generated at the first larval stage (e.g. PVX of the P cell lineage), and they continue to be generated during the L2 stage (e.g. PCB), L3 stage (B and P cell descendants), and L4 stage (mostly ray sensory neurons)(Sulston et al., 1980). Strikingly, we find that despite their distinct generation time, the expression of the more than 39 terminal differentiation markers (located on the NeuroPAL transgene) are tightly coordinated over a relatively small window during the mid-to-late L4 stage (**Fig.7; Suppl. Fig.S2**). At early L4, color codes are not yet established (**Fig.7**), nor are they at earlier larval stages (**Suppl. Fig.S2**). The mid-to-late L4 stage is concomitant with the beginning of male tail retraction, a sexually dimorphic process that results in the generation of male-specific mating organs (Emmons, 2005; Nguyen et al., 1999; Sulston et al., 1980). The extent of this coordination is striking, not only because of the substantial number of cell types over which we observe this coordination, but also because of the breadth of the distinct molecular features that are covered by these molecular markers. Among the components of this marker set are the *cis*-regulatory elements from four distinct panneuronal genes (*unc-11, rgef-1, ehs-1, ric-19*). Like the neuron-class-specific marker genes, all these elements only begin to drive reporter gene expression during the mid-to-late L4 stage, concomitant mail tail retraction.

**Fig.7:**
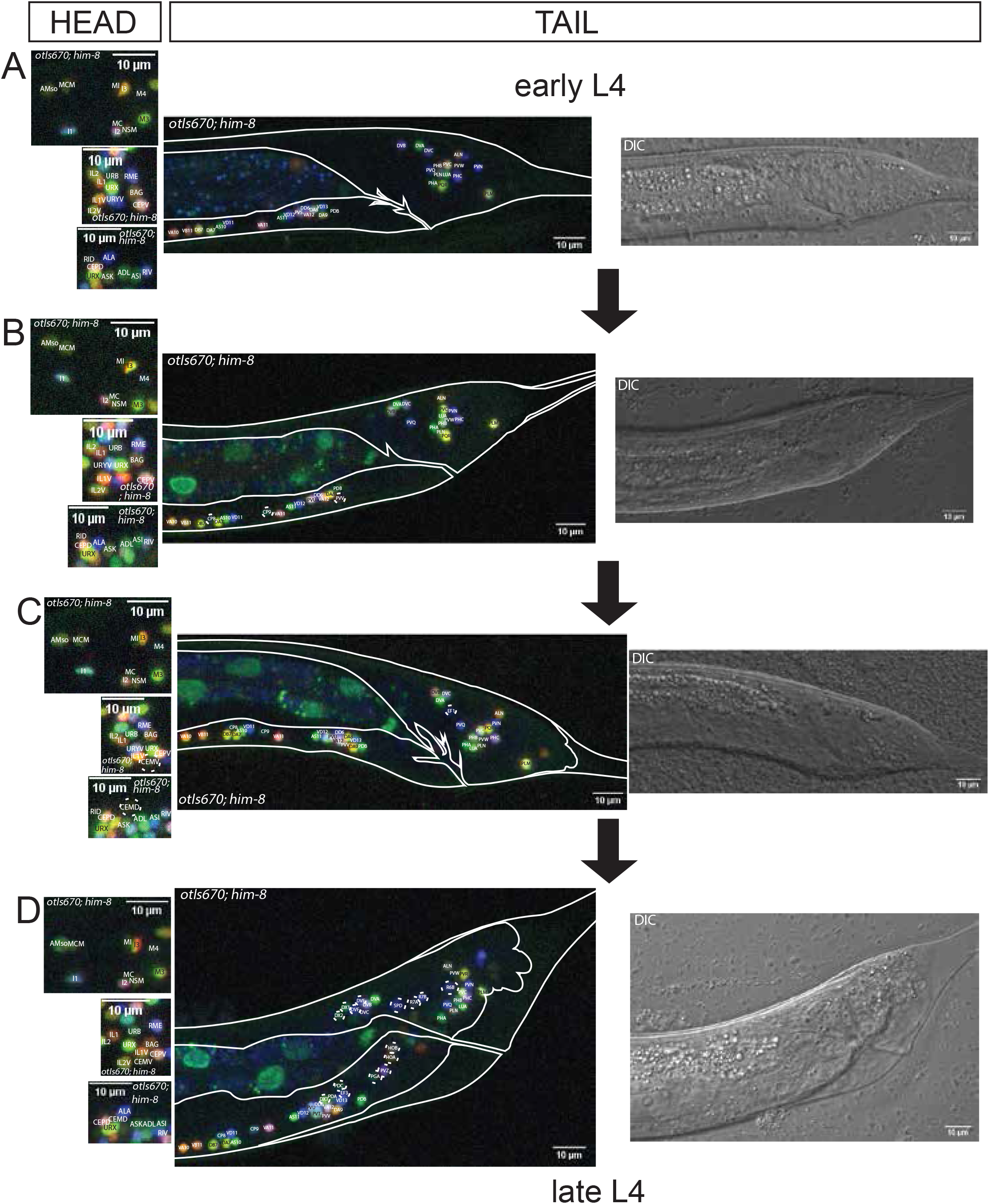
Emergence of the color code in males: L4 substages. Male-specific neuronal color code emergence in the male tail (*otIs670; him-8* genotype) is coordinated with retraction of the tail, which occurs 4 hours before the L4-to-adult molt. The leftmost panel depicts representative images of NeuroPAL expression in the heads of L4 males in the following ganglia: the anterior pharynx, the anterior ganglion, and the dorsal ganglion. The middle panel shows the male NeuroPAL expression from early to late L4. The rightmost panel depicts the DIC image of each stage of the L4 male tail. All images are representative of the 20 worms examined for each of the indicated L4 stages. **A:** A representative image of an early L4 male tail and age-matched images from the head. At this stage only the sex-shared neurons in the tail are present with their NeuroPAL color. **B:** A representative image of an L4 male tail that has just begun to retract and age-matched images from the head. At this stage the male neurons indicated in dashed circles, CP8, CP9, and PVV, are present with their adult color codes. **C:** A representative image of an L4 male tail that has further retracted, and images of age-matched regions in the head. At this stage in the tail, the male neuron EF1 (indicated in the dashed circle) is present with the adult color code. At this L4 stage, the male-specific head neurons CEMV and CEMD neurons (dashed circles) are present with their adult color codes in the anterior and dorsal ganglions, respectively. Thus at this stage all the male-specific neurons in the head have acquired their adult color codes. **D:** Representative image of a late L4 male tail and images of age-matched regions in the head. At this stage in the tail, many of the male-specific neurons have their adult colors, as indicated in the dashed circles.

None of the markers for which we observed a coordinated, delayed onset during the L4 stage in sex-specific neurons, display a delay in sex-shared and embryonically born neurons. That is, all NeuroPAL markers turn on after neuron birth and remain stable throughout all larval stages (**Suppl. Fig.S2**). The best illustration of the specificity of this coordinated expression wave in male-specific neurons are the sex-shared neurons. These neurons are born in both sexes at similar stages of larval development. We consistently find that in all cases, the NeuroPAL color code turns on shortly after the generation of these neurons (*i.e*., after their terminal cell division)(**Suppl. Fig.S2**). For example, the sex-shared PDB and VD13 neurons, both close lineal relatives to the male-specific PVX neuron, are born at about the same time as the male-specific PVX (**Fig.1B**). While PVX generates its color code only at the L4 stage, PDB and VD13 generate their color code in the L1 stage after their birth (**Suppl. Fig.S2**). Similarly, L2-generated, male-specific PCB neuron class turn on their color code at the mid-to-late L4 stage, while the L2-generated, but sex-shared RMF neuron class and the L2-generated neurons of the postdeirid lineage generate their color code shortly after their birth (**Fig.7**). We conclude that earlier born male-specific neurons delay their differentiation until the fourth larval stage to then differentiate in a coordinated manner, concomitantly with the differentiation of sexual organs.

## DISCUSSION

The nervous system of the *C. elegans* male contains almost 30% more neurons than the *C. elegans* hermaphrodite. The male-specific nervous system is structurally complex and controls the many intricate steps of male-mating behavior (Garcia et al., 2001; Liu and Sternberg, 1995). Even behaviors that are not directly related to mating behavior – from chemosensation, exploratory behavior, and locomotion to defecation – are sexually dimorphic but the extent to which the male-specific nervous system is involved in these dimorphisms remains to be explored (Barr et al., 2018; Portman, 2017). From a developmental perspective, it is fascinating to ask how complex interconnected circuitry is established during postembryonic development and integrated with an already existing nervous system. To address the many questions relating to the development and function of the male-specific nervous system, it is of interest to be able to characterize gene expression and gene function within it and also visualize its neuronal activity patterns. The tool that we present here, NeuroPAL, presents a major stepping stone to achieve these goals. NeuroPAL enables rigorous analysis of gene expression patterns. Moreover, the ability to combine NeuroPAL with GCaMP-based neuronal activity recordings – as recently shown in the hermaphrodite nervous system (Yemini et al., 2021) – will pave the way to decipher neuronal activity patterns in the male nervous system.

Owing to its ability to visualize the expression of more than 40 distinct genes that measure the live differentiated state of neurons throughout the entire nervous system of both sexes, we have now been able to use NeuroPAL to gain novel insights into the development of the male-specific nervous system. We corroborated and extended the findings of an unusual non-stereotypic variability in the generation of a specific set of neurons, the EF and DX neurons. We further refined and also revealed the novel patterning roles of three gene regulatory factors: a Hox cluster gene, a miRNA and a bHLH transcription factor. Perhaps most interestingly, we used NeuroPAL to reveal that despite their diverse birth dates, male-specific neurons coordinate the acquisition of terminal identity features to within a specific window of time, the mid-to-late fourth larval stage. At this time other non-neuronal mating structures – including fans, rays and spicules – are generated (Emmons, 2005; Nguyen et al., 1999; Sulston et al., 1980). We term this coordinated differentiation “just-in-time” differentiation, to illustrate that neurons only acquire their functional properties once all the “effector systems” of the male-specific nervous system (*i.e*., all the end organs innervated by the male-specific neurons) come into existence and once the mating process becomes physically possible due to the generation of such mating organs. It is important to emphasize the two reasons why “just-in-time” differentiation cannot merely be a reflection of a delay in maturation of the fluorophores, with which we measure differentiation programs. First, fluorescent signals are visible in sex-shared neurons immediately after their generation at early larval stages, whereas male-specific neurons that were born concurrently show a delay of up to several larval stages (>24 hours later), whereas fluorophore maturation times are known to operate on a much faster scale, with most maturing within <1 hour (Balleza et al., 2018; Cranfill et al., 2016). Second, the birth dates of different male-specific neurons is distinct, yet the onset of fluorophore expression is coordinated to occur at the same time.

The “just-in-time” terminology is adapted from “just-in-time” specific transcriptional programs in metabolic pathways (Zaslaver et al., 2004). Genes that code for specific proteins in the metabolic production machinery display temporal dynamics which ensure that, when a metabolic production pipeline is being ramped up under specific conditions, proteins are generated only when needed in the production pipeline. This allows the machinery to reach a production goal with minimal total enzyme production (Zaslaver et al., 2004).

Coordinated, “just-in-time” differentiation programs are visualized by the NeuroPAL transgene throughout all male-specific neurons. For one of them, the CEM neurons, the delayed onset of differentiation had been previously noted before. The CEM neurons are born in the embryo (and die in hermaphrodites) but were reported to initiate expression of several molecular features, including the putative sensory receptor *pkd-2* and its cholinergic-neurotransmitter phenotype, only by the L4 stage (Lawson et al., 2019; Pereira et al., 2015; Wang et al., 2010). CEM neurons only synapse onto sex-shared neurons that were generated and already differentiated in the embryo, thus the “just-in-time” differentiation of CEMs at the L4 stage cannot simply relate to the appearance of sex-specific effectors cells at the L4 stage. The reason that CEM differentiation is delayed until the L4 stage likely lies in their function: CEM neurons sample mating cues (Narayan et al., 2016; Srinivasan et al., 2008) and hence are not required to operate until the male is sexually mature.

“Just-in-time” differentiation is not unique to male-specific neurons. Hermaphrodite specific neurons, of which there are only two classes, the HSN and VC neuron classes, had already been reported to acquire their fully differentiated state only in the L4 stage. In the HSN neurons, which are embryonically born, this is best evidenced by the acquisition of its serotonergic neurotransmitter identity, which they acquire at the late L4 stage (Desai et al., 1988). In the VC neurons, which are generated by the late first-larval stage, cholinergic marker gene expression only becomes induced in the L4 stage as well (Pereira et al., 2015). The logic of the “just-intime” differentiation of the HSN and VC neurons is evident: they innervate vulval musculature that only becomes generated and properly placed during late-larval development (Sulston and Horvitz, 1977). Similarly, in the male, individual end organs innervated by male-specific neurons (*e.g*., spicule muscle) are only generated and put in place at late larval stages. Moreover, male-specific neurons that are synaptically connected, are generated at distinct larval stages (*e.g*., the PVX neuron, which is born in the first larval stage, synaptically connects with the male-specific PCA, CP, and ray neurons, which are generated at L3 or L4 larval stages). The “uncoordinated” generation of multiple components of the male tail may therefore explain why terminal differentiation of male-specific neurons has to be coordinated.

How is this coordinated, just-in-time differentiation wave genetically specified? For the proper timing of differentiation of the male-specific CEM and hermaphrodite-specific HSN neurons, the heterochronic pathway has been implicated (Lawson et al., 2019; Olsson-Carter and Slack, 2010). This pathway is composed of a series of sequentially activating gene-regulatory factors, including transcription factors, regulatory RNAs and translational regulators (Rougvie and Moss, 2013). However, the effects of this pathway on CEM and HSN timing was shown to be only partial (Lawson et al., 2019; Olsson-Carter and Slack, 2010), suggesting the involvement of other regulatory factors. For example, it can be envisioned that target-derived signals help to coordinate the timing of “just-in-time” differentiation. Perhaps sex-specific neurons are under a “differentiation break” that actively inhibits the execution of terminal differentiation. This notion is again inferred from the CEM and HSN neuron cases, whose identity-specifier, the terminal selector UNC-86 (Lloret-Fernandez et al., 2018; Shaham and Bargmann, 2002), is already present in the CEM and HSN neurons since their birth (Finney and Ruvkun, 1990), but is unable to induce CEM differentiation until the time is right. Correctly-timed induction might thus be achieved by an inhibitory mechanism that prevents UNC-86 function or, alternatively, by the absence of a critical UNC-86 cofactor, whose expression is temporally controlled.

An intrinsic feature of terminal differentiation programs of many, and perhaps all *C. elegans* neurons, may explain why terminal differentiation of sex-specific neurons appears to be an all-or-nothing event. In many, and perhaps all *C. elegans* neurons, gene expression programs within a neuron are highly coordinated via the activity of terminal-selector transcription factors that become active right after the birth of a neuron in order to initiate terminal differentiation (Hobert, 2016). Triggering the entire differentiation program of a neuron prematurely, *i.e*. before needed, and thus not coordinated with the differentiation of other neurons, may send uninformative or even conflicting signals to the sex-shared nervous system. Just-in-time differentiation, coordinated over multiple cell and tissue types, ensures that individual components of a nervous system only go online once every individual component is set in place.

## MATERIAL AND METHODS

### Strains

The following mutant alleles were used in this study: *lin-4(e912), egl-5(u202), and lin-32(tm1446)*. The following reporter strains were used: NeuroPAL strains *otIs669; him-5* and *otIs670; him-8* (Yemini et al., 2021). Reporter strains are *flp-3*(*syb2634[T2A::3xNLS::GFP]*) (generated by Sunybiotech) and *flp-27*(*syb3213[T2A::3xNLS::GFP]*) (generated by Sunybiotech). In both cases, the reporter cassette was inserted at the 3’ end of the gene.

### Microscopy

Worms were anaesthetized using 20 mM sodium azide and were mounted on 5% agarose pads. Images were acquired on a Zeiss LSM880 confocal microscope, equipped with 7 laser lines: 405, 458, 488, 514, 561, 594, and 633 nm and processed using ImageJ software. Gamma was adjusted for maximal color distinction for all images. All reporter and mutant strains were imaged at 40X. To obtain worms staged throughout larval development, adult hermaphrodites were allowed to lay eggs for one hour. After one hour, the hermaphrodites were removed from the plates. The plates were then stored at 20°C until the time points corresponding to each larval stage when they were imaged.

#### Variability in cell position

To generate the statistical atlases of male neuron positions, their variability, and colors, we used our previously published strategy (Yemini et al., 2021). Briefly, we initialized the atlas to be the point cloud of one of the worms, then iteratively aligned each animal’s neuron position point cloud to the current atlas, until all animals were aligned. These point clouds, which represent neuron positions and colors, were extracted from images of 12 male animals that were manually annotated. Males were age-matched to those used in our previously published hermaphrodite atlas (Yemini et al., 2021). For each neuron, the atlas includes a mean position, mean color, and a covariance matrix representing the variability for the neuron’s position and color across the population of worms. We computed the positions of the male and hermaphrodite neurons using the determinant of the neuron’s positional covariance matrix as an estimate of their aligned spatial occupancy volume (**Table S2**). We used these volumes to compute differences in neuron position variability between hermaphrodites and males. To account for color variability resulting from any changes in the imaging hardware (*e.g*., aging equipment such as excitation lasers) we affine transformed the male atlas colors to those of the hermaphrodite using the sex-shared neurons. Color alignment and histogram matching have been previously shown to be a critical step for downstream analysis in such computations as atlas creation and neural segmentation (Nejatbakhsh et al., 2020; Varol et al., 2020).

## ACKNOWLEDGEMENTS

We thank Rene Garcia for help in distinguishing DVE and DVF, Laura Peirera for help in generating the male NeuroPAL map, and members of the Hobert lab for comments on the manuscript. This work was funded by the NIH (R37NS039996) and the Howard Hughes Medical Institute.

## SUPPLEMENTARY LEGENDS

**Table S1: List of all male-specific neurons and their color signal in NeuroPAL.**

Because different promoters drive the same reporter fluorophores, the assignment of cell-specific color codes relies to a large extent on the examination of each reporter on its own. In the context of the male-specific nervous system, this has been done for some, but not all of these reporters. Wherever we have confirmed expression of the promoter in a neuron, the box is filled with the color corresponding to its reporter-fluorophore origin. Wherever we cannot be sure from where the color in a particular neuron originates, the box is filled with gray. Wherever we are sure there is no expression in a neuron, the box is left white.

**Table S2: Positional variability and color of all neurons in males and hermaphrodites**

The positional and color variability of each neuron in the hermaphrodite and male head and tail. The results are based on 10 hermaphrodites, 12 male heads, and 13 male tails.

**Suppl. Fig.S1: Positional variability in the male versus hermaphrodite head**

**A:** Ventral view of the positions of the hermaphrodite (red) and male (blue) neurons (circles) in the tail. The sex-shared neurons are linearly aligned to each other, labeled, and corresponding pairs are connected by a line. Note that, whereas most sex-shared neurons are positioned similarly in both sexes, the beginning of the hermaphrodite VNC is displaced anterior to its male counterpart and, in contrast, the hermaphrodite dorsorectal ganglion neurons are displaced posterior to their male counterparts.

**B-D:** The atlas of male neuron positional variability (based on 12 male heads) for the left (B), right (C), and ventral (D) sided views. Dots indicate the mean position of each neuron. Ellipses indicate the positional variability of each neuron in the given axis. Neurons colors approximate those in NeuroPAL but have been brightened for visibility.

**Suppl. Fig.S2: NeuroPAL in male head and tail L2-L3**

Representative images of NeuroPAL expression (*otIs670;him*-8) in larval stage males (early L2, late L2, L3). The leftmost panel depicts representative images of NeuroPAL expression in the heads of larval L4 males in the following ganglia: the anterior pharynx, the anterior ganglion, and the dorsal ganglion. These ganglia represent the regions where male-specific neurons are located in the adult male head. The rightmost panel depicts representative images of NeuroPAL expression in the tails of larval L4 males.

